# Synthesis of Long RNA with a Site-Specific Modification by Enzymatic Splint Ligation

**DOI:** 10.1101/2022.09.17.508400

**Authors:** Howard Gamper, Caroline McCormick, Sepideh Tavakoli, Meni Wanunu, Sara H. Rouhanifard, Ya-Ming Hou

## Abstract

Synthesis of RNA molecules that contain an internal site-specific modification is important for RNA research and therapeutics. While solid-state synthesis is attainable for such RNA in the range of 100 nucleotides (nts), it is currently impossible with kilobase (kb)-long RNA. Instead, long RNA with an internal modification is usually assembled in an enzymatic 3-part splint ligation to join a short RNA oligonucleotide, containing the site-specific modification, with both a left-arm and a right-arm long RNA that are synthesized by *in vitro* transcription. However, long RNAs have structural heterogeneity and those synthesized by *in vitro* transcription have 3’-end sequence heterogeneity, which together substantially reduce the yield of 3-part splint ligation. Here we describe a method of 3-part splint ligation with an enhanced efficiency utilizing a ribozyme cleavage reaction to address the 3’-end sequence heterogeneity and involving DNA disruptors proximal to the ligation sites to address the structural heterogeneity. The yields of the synthesized kb-long RNA are sufficiently high to afford purification to homogeneity for practical RNA research. We also verify the sequence accuracy at each ligation junction by nanopore sequencing.

## INTRODUCTION

Synthesis of kb-long RNA with an internal modification is important for probing the structure and function of the epitranscriptome. Matured mammalian mRNAs are now known to contain post-transcriptional modifications (1,2). While each modification in mRNA occurs at a much lower frequency relative to tRNA and rRNA, each may confer information that regulates gene expression at a complexity that opens a new avenue of research (3–5). A critical barrier to progress is the lack of a robust method that precisely maps the position of each mRNA modification in the epitranscriptome. For example, the modification pseudouridine (ψ) in mRNA (6) does not affect Watson-Crick base pairing and thus cannot be detected by hybridization-based sequencing methods, such as by reverse-transcription of RNA during cDNA synthesis for Illumina sequencing. Importantly, nanopore sequencing of RNA has emerged as an attractive alternative (7). In this technology, native mRNA is fed directly into the flow cell without the need to convert into cDNA. Detection of ion current differences for different k-mer sequences (k = 5) during translocation of the RNA through the pore suggests the presence of a modified nucleotide (8–12). However, while the nanopore platform has the unique advantage of producing direct and long reads in a high-throughput capacity, each modification is read as basecalling “mismatch” errors (9,10,13–15). The significance value of each mismatch error is dependent on the sequence context, due to the presence of RNA intramolecular structures that can influence the kinetics of RNA translocation through the pore (16). To quantify the potential of using the basecalling error as an indicator for a modification, the technology needs a synthetic mRNA control with the modification at its homogeneity (i.e., 100%) and in its natural sequence context. This synthetic control is a necessary reference to determine the level of detection by the basecalling error in nanopore sequencing. Additionally, besides contributing to nanopore RNA sequencing technology, long RNA with an internal modification is of great interest to current efforts in RNA therapeutics and vaccine development (17,18).

Synthesis of short RNA (< 100 nts) with an internal modification is achievable through the solid-phase platform of chemical coupling (19–22). This platform, however, is expensive and has a steep decline in product yield with increasing RNA length. In a more cost-effective approach, short RNA with a modification can be synthesized by a 2-part splint ligation that joins two RNAs, one of which contains the modification, on a complementary single-stranded DNA splint (23) (Figure 1A). If the two arms share complementary sequences, such as those that constitute a fulllength tRNA, the enzymatic joining can be facile even without a splint (23). The joining of two single-stranded RNAs in the absence of a splint is preferred by T4 RNA ligase 1 (RNL1) (24), whereas the joining of a nicked RNA in the presence of a splint is preferred by T4 RNA ligase 2 (RNL2) (25,26). The joining can also be achieved by 3-part ligation (Figure 1B), where the middle RNA is synthesized by a solid-phase approach with the site-specific modification, whereas the two side RNAs are each made by *in vitro* transcription, usually with T7 RNA polymerase (RNAP) (23,27). The three RNAs are aligned on a DNA splint and are joined by RNL2. For assembly and synthesis of a short RNA, the efficiency of 2-part or 3-part ligation is typically 30-60%, which is a practical yield that affords purification and subsequent analysis. Even if the short RNA has a stable structure, such as the well-defined structure of a tRNA, the ligation position can be chosen to minimize structural interference (23).

**Figure 1.**
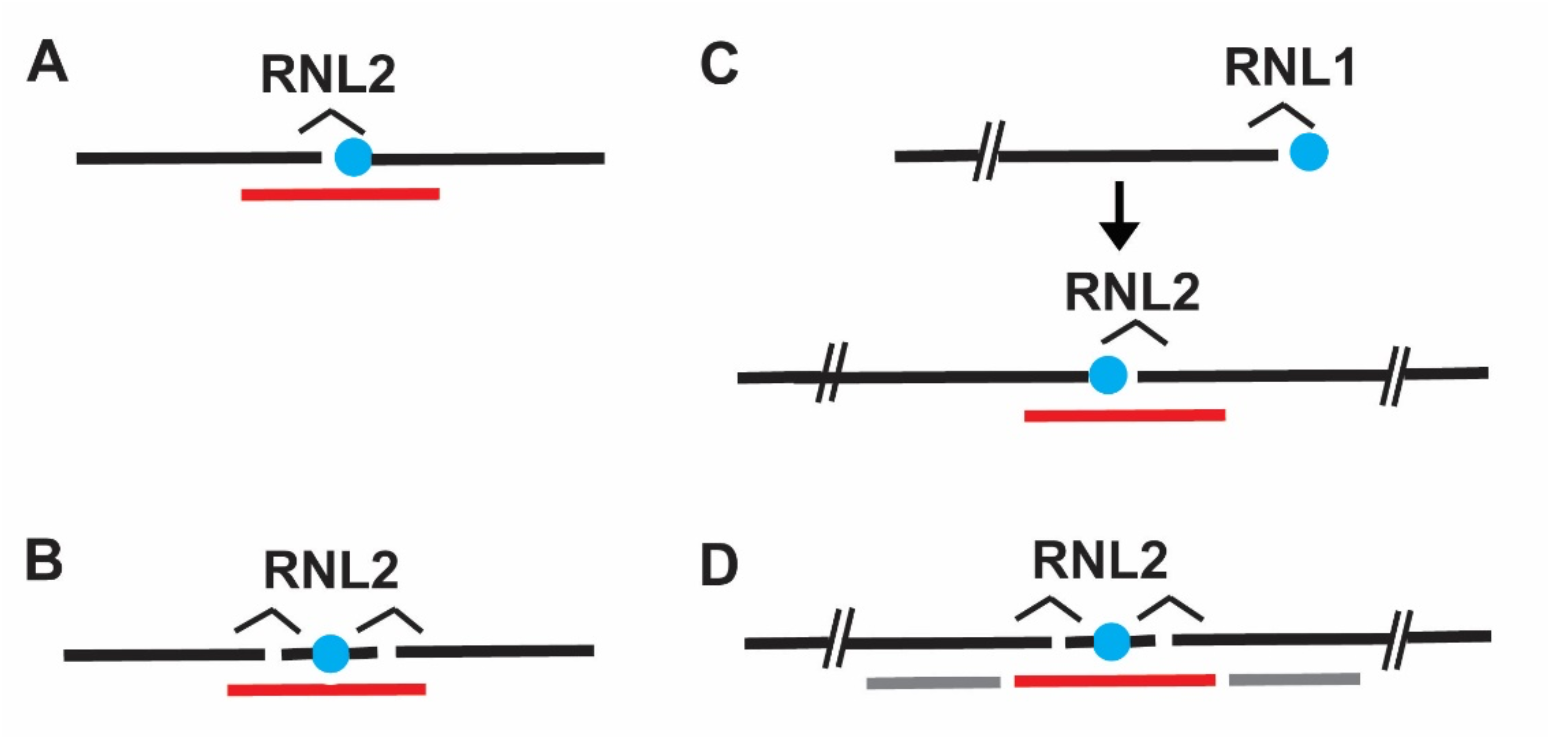
Four methods of assembly to synthesize RNA containing a site-specific internal modification. (A) Assembly of a short left-arm RNA with a short right-arm RNA, the latter of which has a 5’-terminal modification in a 2-part splint ligation. (B) Assembly of a short left-arm and a short right-arm RNA with a modification-containing middle RNA in a 3-part splint ligation. (C) Terminal 3’-extension of a long left-arm RNA with a modified nucleoside 3’,5’-bisphosphate, followed by removal of the 3’-phosphate by an alkaline phosphatase, and joining with a long rightarm RNA in a 2-part splint ligation. (D) Assembly of a long left-arm RNA and a long right-arm RNA with a short middle RNA with the internal modification in a 3-part splint ligation, in the presence of both a left-arm and a right-arm DNA disruptor. Short RNA (less than a 100-mer) is shown as a straight black line, whereas long RNA (more than a 100-mer) is shown as a straight black line with double daggers. The modified nucleotide is shown as a cyan dot, the splint DNA is shown in red, and the DNA disruptors are shown in grey.

In contrast, kb-long RNA cannot be generated in full-length by solid-phase synthesis. The current technology of solid-phase synthesis is limited to fewer than 200 nts but with poor yield and frequent synthesis failure. Instead, kb-long RNA must be assembled from fragments by a combination of enzymatic and chemical synthesis. One such method employs RNL1-dependent extension of an *in vitro* transcribed left-arm RNA with a modified nucleotide, which is then joined by an RNL2-mediated splint ligation with the right-arm RNA, also generated by *in vitro* transcription (Figure 1C) (28). In this method, the modified nucleotide is synthesized with 3’,5’-bisphosphates, which restrict extension of the left-arm to a single nucleotide using the 3’-phosphate as a block (29). After dephosphorylation of the 3’-phosphate, the extended left-arm is joined with the right-arm by a 2-part splint ligation. While this method successfully generated a ~400-mer RNA (28), synthesis of the modified nucleotide with 3’,5’-bisphosphates requires special expertise. A more general method is a 3-part splint ligation to join a chemically synthesized short RNA that bears the site-specific modification with a left-arm and a right-arm RNA, the latter of which are synthesized by *in vitro* transcription. However, the yields have been low (< 2%) (30), preventing practical purification and subsequent study of the ligated RNA.

Here we report a method that improves the yield of 3-part ligation to synthesize kb-long RNA containing a site-specific internal modification. The innovation is the combination of two strategies that address key issues of low yield. First, *in vitro* transcribed RNA usually has a population of 3’-ends, due to the propensity of RNAPs to prematurely terminate, and alternatively, to extend beyond the 3’-end with extra non-templated nucleotides (31–33). This problem was previously addressed on short RNA by transcribing the RNA with a *cis*-acting ribozyme that would self-cleave, leaving the dissociated short RNA with a homogeneous 3’-end (34,35). Second, long RNA has inherent structural heterogeneity, which lacks a well-defined tertiary structure but folds and re-folds spontaneously and dynamically with the ability to engage both termini in intramolecular base pairing, thus blocking them from splint ligation. It was found that using DNA disruptors to hybridize to RNA sequences near each ligation site or using an ultra-long DNA splint (up to 100 nts) improves the ligation yield (30,36,37), presumably by freeing up the RNA termini. We have incorporated both strategies into a single method that improves the ligation yield by 3-5-fold over the best reported yields (30,37). In this method, we engineer a *cis*-acting ribozyme to the left-arm RNA to produce a precise 3’-end, and we include two DNA disruptors to hybridize next to the ligation sites in the ligation reaction. In addition, we describe for the first time a method to purify 1 kb-long RNA for sequence verification of ligation accuracy, using nanopore sequencing at single-molecule resolution. Combined, this method demonstrates the ability to generate kb-long RNA bearing a site-specific modification for broader research.

## MATERIALS AND METHODS

### Synthesis of left-arm and right-arm RNAs by *in vitro* transcription

The template for *in vitro* transcription for synthesis of a left-arm or a right-arm RNA, each ~500-mer and unmodified, is made by solid-phase synthesis as a double-stranded gBlock DNA by IDT (Integrated DNA Technologies). A gBlock double-stranded DNA is designed with the consensus T7 promoter sequence, followed by the sequence of interest beginning with three G nucleotides to facilitate transcription. The gBlock for the left-arm RNA additionally encodes the sequence for the Hepatitis delta virus (HDV) ribozyme (38,39), which upon synthesis by transcription can self-cleave to release the left-arm RNA. The HDV ribozyme has the sequence: 5’-GGGUCGGCAUGGCAUCUCCACCUCCUCGCGGUCCGACCUGGGCUACUUCGGUAG-GCUAAGGGAGAAG-3’

Left-arm and right-arm gBlocks (500 ng, ~1.5 pmoles) were transcribed at 37 °C, 2h, in 20 μL using the NEB HiScribe kit. Because the right-arm RNA must have a 5’-monophosphate (5’-pG) to participate in the ligation reaction, its transcription reaction was supplemented with 20 mM GMP and 10 mM MgCl_2_. After transcription, the gBlock DNAs were hydrolyzed by RNase-free DNase I (NEB) at 37 °C, 15 min, and the RNA products were isolated using the NEB 50 μg-scale Monarch RNA Cleanup cartridges. The yield of purified RNA is usually > 100 μg.

Each *in vitro* transcribed RNA was determined for the concentration by A_260_ (usually in a 1:50 dilution of the stock) and analyzed (usually 1 μL of the 1:50 dilution) in a 6% denaturing PAGE/7M urea gel (abbreviated as denaturing PAGE hereafter). The gel was run in 1X TBE (90 mM Tris, pH 8.0, 90 mM boric acid, 2 mM Na2EDTA) in a Bio-Rad mini-Protean apparatus for 30-60 min at 200 V at 60 °C, along with a low MW (molecular weight)-range RNA ladder (NEB). SYBR Gold-stained gels were imaged to determine the fraction of intact RNA in each sample, which was used to adjust the RNA concentration as determined by A_260_ to more accurately reflect the concentration that would participate in the ligation reaction. Additional assessment of the RNA concentration was obtained by comparing the RNA band intensity to the known amount of a standard RNA in the same gel.

### Ribozyme self-cleavage of the left-arm RNA to generate a homogeneous 3’-end

While HDV can catalyze self-cleavage during transcription, this activity is usually at a low level. To produce a higher level of cleavage for better ligation yield, several heat-cool cycles were performed. The *in vitro* transcribed left-arm RNA (200 pmoles) was mixed with a 60-mer left-arm DNA disrupter (2 nmoles) in 90 μL of 110 mM Tris-OAc (Tris acetate, pH 6.3). The reaction was incubated at 85 °C, 2 min, cooled to room temperature, and supplemented with 5 μL 200 mM MgCl_2_ and 5 μL 200 mM 2-mercaptoethanol (β-Me) to a final volume of 100 μL (at the final concentration of 10 mM MgCl_2_, 10 mM β-Me, 20 μM of the left-arm DNA disruptor, and 2 μM of the left-arm RNA). The reaction was transferred to a PCR tube and incubated in a thermocycler at 72 °C, 30 s, followed by 4 heat-cool cycles each lasting 15 min between 72 °C and 8 °C. The yield of the cleavage was determined by the fraction of the cleaved product in the total input leftarm RNA. The A_260_ was not informative, due to the presence of disruptor DNA.

### Hydrolysis of 2’,3’-cyclic phosphate on the left-arm RNA generated by HDV cleavage

HDV cleavage of the transcribed left-arm RNA produces a 2’,3’-cyclic phosphate at the 3’-end (40), which was hydrolyzed by adding 1.5 μL of 10 units/μL T4 PNK (polynucleotide kinase) and 1 μL RNase-Out solution (40 units/μL, ThermoFisher) to the cleavage reaction above. The hydrolysis reaction was incubated at 37 °C, 8 h, while aliquots of 0.4 μL were analyzed on a 6% analytical denaturing PAGE. Each reaction aliquot as well as a final reaction mixture was extracted with phenol:chloroform:isoamyl alcohol (25:24:1), pH 5, and the RNA was ethanol precipitated with 1/10 vol of 2.5 M NaOAc, pH 5.0, and 3 vol ethanol. The pellet was washed, air dried, and dissolved in 20 μL RNase-free water. Hydrolysis was verified by ligation of the T4 PNK-treated left-arm RNA with a 15-mer RNA in a 2-part ligation reaction, using the same splint and conditions as in the 3-part ligation reaction.

### Design of DNA disruptors for 3-part splint ligation

DNA 60-mer disruptors were designed to hybridize to the left-arm and right-arm RNA adjacent to the 3’- and 5’-end of the DNA splint in each 3-part splint ligation reaction (see Figure 2A). These DNA disruptors were synthesized by IDT without purification.

**Figure 2.**
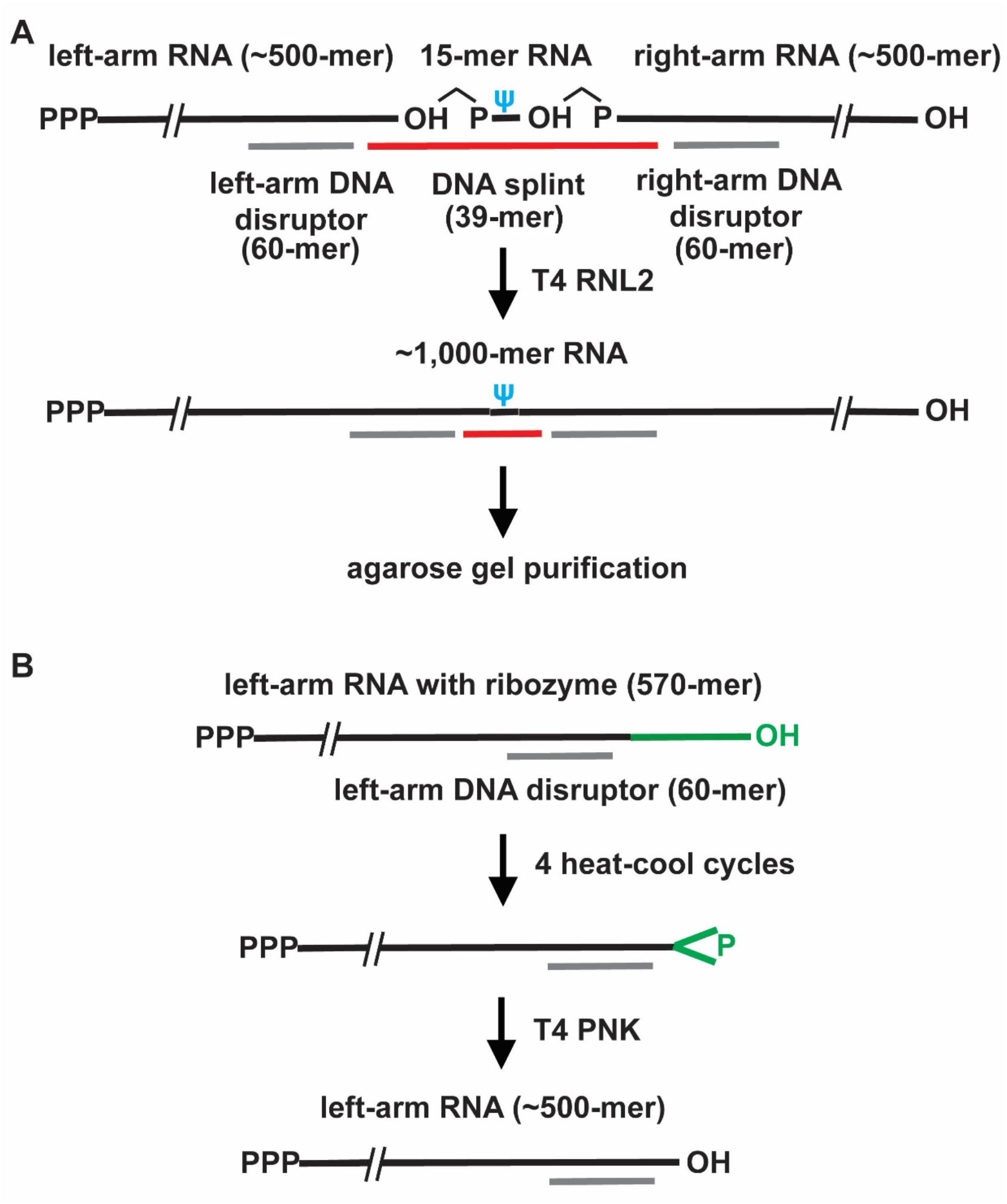
Scheme of the 3-part splint ligation of this work. (A) The matured left-arm RNA (~500-mer) is transcribed with a 5’-triphosphate and is processed by HDV and T4 PNK to produce a homogeneous 3’-OH, the matured right-arm RNA (~500-mer) is transcribed with a 5’-monophosphate, while the middle RNA (15-mer) containing a site-specific internal ψ is chemically synthesized. The three RNAs are assembled on a DNA splint (39-mer), in the presence of a leftarm and a right-arm DNA disruptor (60-mer each), for joining by T4 RNL2 to produce an ~1kb RNA. (B) The left-arm RNA is transcribed as a fusion with the HDV ribozyme (in green) to produce a 570-mer, in which HDV self-cleaves to release the left-arm RNA (~500-mer) with a 2’,3’-cyclic phosphate. Treatment with T4 PNK generates the mature 3’-OH end. In each sequence context, the left-arm and the right-arm RNA are synthesized in the range of a 500-mer.

### Three-part splint ligation

The molar ratios below provide a framework to optimize each reaction. A typical 3-part ligation reaction consists of 1:1:1.5 ratio of the left-arm RNA, the right-arm RNA, and the 15-mer RNA that is chemically synthesized with a site-specific modification. These RNAs are then mixed with a 24:24:0.9 molar ratio of the left-arm DNA disruptor (60-mer), the right-arm DNA disruptor (60-mer), and the DNA splint (39-mer). For the reactions described here, the molar ratios represent 15 pmoles of the ribozyme-cleaved and T4 PNK-treated left-arm RNA, 15 pmoles of the right-arm RNA, 22.5 pmoles of the 15-mer RNA with a modification, 360 pmoles of the left-arm DNA disruptor, 360 pmoles of the right-arm DNA disruptor, and 13.5 pmoles of the splint DNA.

All components of a 3-part splint ligation reaction were mixed in a PCR tube according to the molar ratios above with 3.6 μL of 0.5 M HEPES, pH 7.5, in a total volume of 27 μL. The 3-part ligation reaction was initiated at 40 °C, 5 min, and cycled down to 5 °C by decreasing 5 °C every 2 min. The annealed 3-part pre-ligation complex was mixed with 9 μL of a 4X ligation sub-stock to 36 μL and incubated at 16 °C, 15 min, 25 °C, 30 min, and 37 °C, 60 min, in a thermocycler. The 4X ligation sub-stock contains 8 mM MgCl_2_, 2 mM ATP/Mg^2+^, 4 mM DTT, 14 μM RNL2, and 2 units/μL RNase-Out. Each 3-part splint ligation was performed in 36 μL with the final concentration of 0.42 μM 3’-end processed left-arm RNA, 0.42 μM right-arm RNA, 0.62 μM 15-mer RNA with a site-specific modification, 10 μM each of the left-arm and right-arm disruptors, 0.38 μM splint DNA, 50 mM HEPES, pH 7.5, 2 mM MgCl_2_, 0.5 mM ATP/Mg^2+^, 1.0 mM DTT, 3.5 μM RNL2, and 0.5 units/μL RNase-Out.

### Clean-up of the 3-part splint ligation reaction

Once completed, the 3-part ligation reaction was diluted to 150 μL with RNase-free water, supplemented with 15 μL 2.5 M NaOAc, pH 5.0, 1 μL 20 μg/μL glycogen, and extracted twice with equal volumes of phenol:chloroform:isoamyl alcohol (25:24:1), pH 5.2. Following an ethanol precipitation, the nucleic acid pellet was dissolved in 15 μL 70% deionized formamide by heating at 65 °C, 5 min. To determine the efficiency of 3-part vs. 2-part ligation, an aliquot of 0.6 μL was run on a 6% denaturing PAGE. Typical yields are 10-35% for 3-part ligation and 35-65% for 2-part ligation.

### Gel purification of the ligated 1 kb RNA for nanopore sequencing

Ligation workups from the previous step were supplemented with 3 μL of 6X purple gel loading dye (NEB), heat denatured at 85 °C for 1 min, and electrophoresed at 100 V, 1 h, in an 1.2% agarose gel (8 x 7 cm) with 6 wells in TAE buffer (40 mM Tris-acetate, pH 8.3, 1 mM EDTA) (41). An authentic 1 kb RNA standard was included as a reference. The ethidium bromide-stained gel was visualized on a BioRad ChemiDoc imaging system, and a paper printout of the image was used as a guide to excise the 1 kb bands of interest. From the agarose gel, 40-50% of the input 1 kb RNA (usually 160-250 ng) was recovered intact in 15 μL water using the Zymo Clean RNA gel recovery kit. Concentration of the RNA was determined using the Qubit RNA HS assay kit and its integrity was assessed by the Agilent 2100 Bioanalyzer with an RNA Nanoreagent chip. Gel-purified 1 kb RNAs from 3-part ligation were combined for Direct RNA sequencing (SQK-RNA002) with the ONT direct RNA sequencing protocol (DRCE_9080_v2_revH_14Aug2019) as described (42). Basecalling, alignment, and signal intensity extraction was as described (42).

### Affinity purification of the ligated 1 kb RNA

Each 3-part splint ligation reaction (36 μL) was extracted twice with phenol:chloroform:isoamyl alcohol (25:24:1), pH 5.0, followed by ethanol precipitation or cleanup through a Zymo RNA Clean and Concentrator-5 cartridge. The recovered RNA, consisting of the 1 kb full-length, the left-arm and right-arm 500-mers, and the 15-mer, was supplemented with 75 pmoles of the biotinylated 39-mer splint and 750 pmoles each of the left-arm and right-arm disruptors in 80 μL of gel elution buffer (0.1% SDS, 1 mM EDTA, 0.3 mM NaOAc, pH 5.0). This solution was carried through a slow heat-cool from 65 °C to 5 °C. It was then added to 40 μL of washed streptavidin-sepharose resin (Cytiva) in gel elution buffer and the mixture was incubated 30-60 min with 1,500 rpm rotation at room temperature. After a brief centrifuge of the suspension at 7,500 rpm, the supernatant was removed. The remaining resin was washed 3 times with 200 μL each of the gel elution buffer and once with TE (10 mM Tris-HCl, pH 8.0). The washed resin was resuspended in 200 μL of fresh TE, and the bound RNA was released to the supernatant by heating at 65 °C, 5 min. After a brief spin, the RNA in the supernatant was precipitated with isopropanol in the presence of 20 μg glycogen, washed, and air dried.

## RESULTS

### Design of a 3-part splint ligation scheme to assemble long RNA

We chose the 3-part splint ligation as a practical method to synthesize kb-long RNA containing a site-specific internal modification. This method is cost-effective, using *in vitro* transcription to synthesize the long left- and right-arm RNA, while using chemical synthesis to generate a short RNA that contains the modification. Because the left-arm and right-arm RNA are transcribed from double-stranded gBlock DNAs, which have the capacity to reach 3 kb, this method in principle can assemble an RNA up to 6 kb long. Additionally, because chemical synthesis can accommodate a wide range of modifications, virtually all naturally occurring mRNA modifications can be addressed. However, current methods of generating long RNA by 3-part splint ligation have low yields (<2%), due to the 3’-end sequence heterogeneity of *in vitro* transcribed RNA, and due to the conformational heterogeneity of long RNA that can shield the termini from ligation. Here we address these obstacles in a 3-part splint ligation method that improves the yield.

We describe the salient features of our method (Figure 2A). (i) The left-arm and the right-arm RNA are synthesized by *in vitro* transcription in the range of a 500-mer, while the middle RNA is chemically synthesized as a 15-mer with the modification in the center separated from the left- and right-end of ligation by 7 nts each. The length of the 15-mer was chosen to maximize the synthesis yield without purification while providing sufficient sequence for splint ligation. The joining of the three RNAs via a splint ligation leads to a product of ~1 kb-RNA, which is suitable for nanopore sequencing to determine the sequence accuracy of ligation and to study the basecalling properties of the modified base. (ii) To facilitate ligation, the right-arm RNA is synthesized with a 5’-p by adding GMP into the NTP mix of *in vitro* transcription. T7 RNAP preferentially initiates RNA synthesis with GMP when it is a component of the reaction mixture (43). The 5’-end of the left-arm RNA is less of a concern for ligation and can be made as a 5’-triphosphate. (iii) To minimize the 3’-end sequence heterogeneity, the left-arm RNA is prepared in two steps (Figure 2B). It is first transcribed with a 3’-end extension to include the HDV ribozyme, which after synthesis catalyzes self-cleavage to release the left-arm RNA with a 2’,3’-cyclic phosphate at the 3’-end. The cyclic phosphate is then hydrolyzed by T4 PNK to generate a homogeneous 3’-end. We choose the HDV ribozyme, which does not have sequence requirements and is broadly applicable to all RNA substrates (44,45). (iv) To minimize the conformational heterogeneity of the left- and right-arm RNA, each is provided with a 60-mer DNA disruptor with a complementary sequence that can hybridize adjacent to the left- and right-end of ligation. The length of the disruptor at 60-mer has been shown to promote assembly of long RNA by 3-part ligation (30). The hybrids are designed to make the termini of the left-arm and right-arm RNA accessible to ligation. While we can also use a long DNA splint, synthesis of DNA of > 100 nts is more expensive, while the ligation yield decreases (37). (v) The ligation reaction is analyzed by a denaturing PAGE, while the full-length 1 kb RNA is extracted from an agarose gel and purified by a Zymo cartridge, which is much easier for long RNA than electro-elution as described in a recent method (37).

We focused on the RNA modification ψ, which is one of the most abundant post-transcriptional modifications in human transcriptome with a frequency of 0.2-0.6% of total uridines (6). RNA modifications with ψ confer resistance to degradation (46) and modulate cellular activities of immunogenicity (47) and translation (48,49). While ψ has been detected by chemical labeling and next-generation sequencing (6,50,51), different labeling methods identify different sites with limited overlap (52). Nanopore sequencing instead has consistently reported it as a U-to-C basecalling mismatch (10,12,53). To quantify the potential of the U-to-C mismatch as an indicator for ψ, we used our 3-part splint ligation method to construct 5 synthetic mRNAs, each bearing a ψ in its natural sequence context in the human transcriptome. Two of these (*PSMB2*; chr1:35603333 (6,51,54) and *MCM5;* chr22:35424407 (6,54)) were annotated with a ψ by previous methods, while the other three (*MRPS14* chr1:175014468, *PRPSAP1*; chr17: 76311411, and *PTTG1P*; chr21:44849705) were detected *de novo* by the U-to-C error in a recent nanopore sequencing analysis (42,55). These 5 synthetic RNAs represent a range of sequence contexts to evaluate our method of 3-part splint ligation.

### HDV cleavage of the left-arm RNA

HDV catalyzes self-cleavage to release the transcribed left-arm RNA with a precise 3’-end (35). We showed that this self-cleavage with long RNA is most effective in multiple cycles of a heat-cool process and, unexpectedly, in the presence of the left-arm DNA disruptor (Figure 3A). With the *MCM5* RNA as an example, the heat-cool cycling alone did not improve the cleavage yield, whereas cycling in the presence of the disruptor did, increasing the yield from 21 to 74% in the 3^rd^ and 4^th^ cycles. Thus, as an RNA itself of a defined tertiary structure (56), the HDV ribozyme (67-mer) refolds with the left-arm RNA most efficiently into the active structure by repeated heat-cool cycles in the presence of the DNA disruptor. This demonstrates the importance of the disruptor to free up the 3’-end of the left-arm RNA for cleavage. Analysis of cleavage of additional left-arm RNAs for *MCM5, MRPS14, PRPSAP1, PSMB2*, and *PTTG1P* supports the importance of the disruptor, showing an increased cleavage yield to 70-90% (Figure 3B).

**Figure 3.**
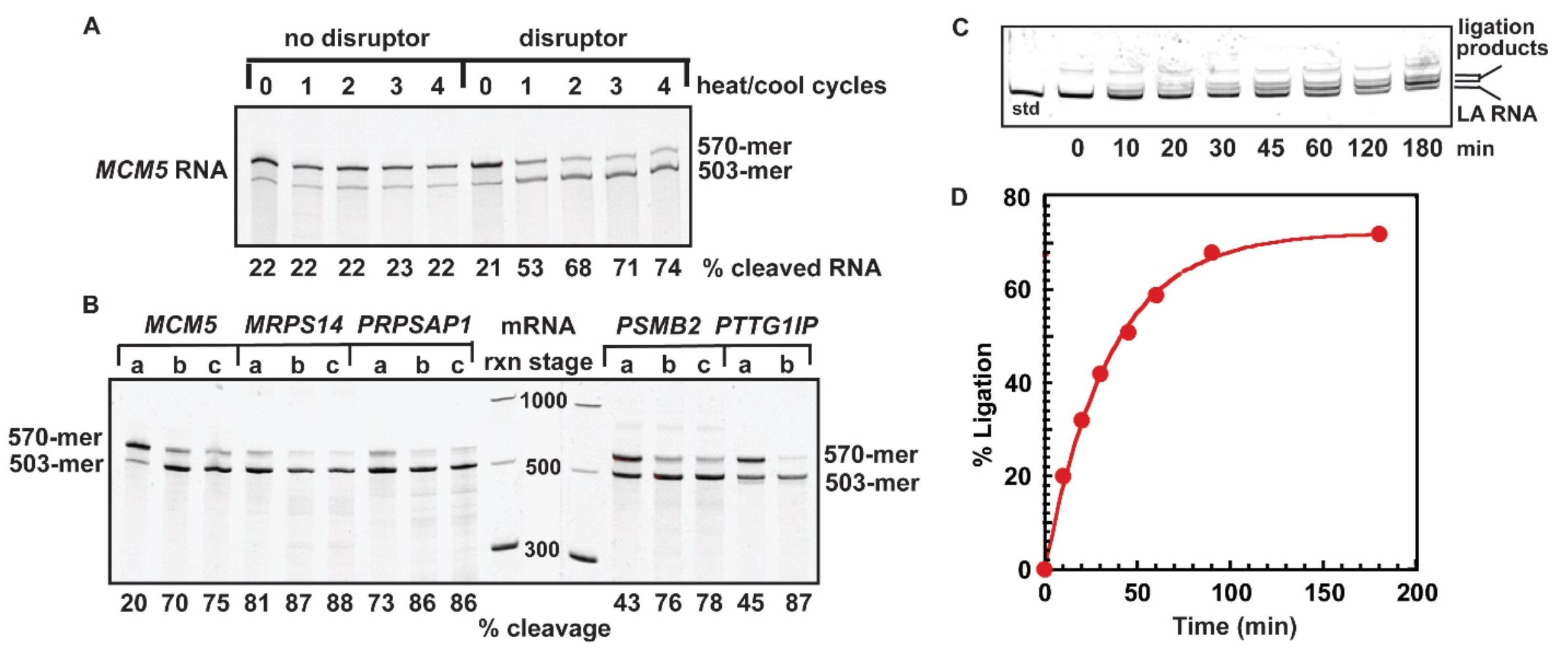
HDV processing of the transcribed left-arm RNA at the 3’-end. (A) Denaturing PAGE (6%) analysis of HDV cleavage of the transcribed left-arm RNA of *MCM5* over the cycling number of a heat-cool process. The left panel was cleavage performed without the left-arm disruptor, while the right panel was cleavage performed with the disruptor, each showing separation of the transcribed (570-mer) from the cleaved RNA (503-mer). The fraction of cleavage was calculated as the band intensity of the 503-mer over the sum of band intensity of the 503-mer and 570-mer. (B) Denaturing PAGE (6%) analysis of cleavage of several transcribed left-arm RNAs, each in three conditions. Conditions “a” denotes no separate incubation for HDV cleavage, “b” denotes HDV cleavage in the presence of the respective left-arm disruptor over 4 cycles of heat-cool, and “c” denotes HDV cleavage and T4 PNK treatment of the HDV-cleaved 3’-end. The fraction of processing by HDV and T4 PNK is shown at the bottom of each condition. (C) Denaturing PAGE (6%) analysis of ligation of the T4 PNK-treated HDV-cleaved left-arm RNA (*PSMB2)* with a 15-mer RNA in a 2-part splint ligation reaction as a function of time of T4 PNK hydrolysis. (D) Efficiency of ligation as measured from data in (C) over time.

HDV cleavage produces a 2’,3’-cyclic phosphate at the RNA 3’-end (40), which needs to be removed before ligation. We used the 3’-phosphatase activity of T4 PNK to hydrolyze the cyclic phosphate and to remove the monophosphate. This T4 PNK reaction did not alter the overall size of each left-arm RNA (Figure 2B), supporting the notion that it is limited to the terminal ribose. The left-arm RNA after T4 PNK hydrolysis was able to join with a 15-mer RNA in a 2-part splint ligation, confirming restoration of the terminal 3’-OH (Figure 3C). In less than 2 h, the ligation efficiency reached a plateau at 70% (Figure 3D), which is the maximum efficiency of 2-part ligation (Figure 5A, below). This indicates that T4 PNK hydrolysis of the cyclic phosphate was stochiometric and completed in 2 h.

**Figure 4.**
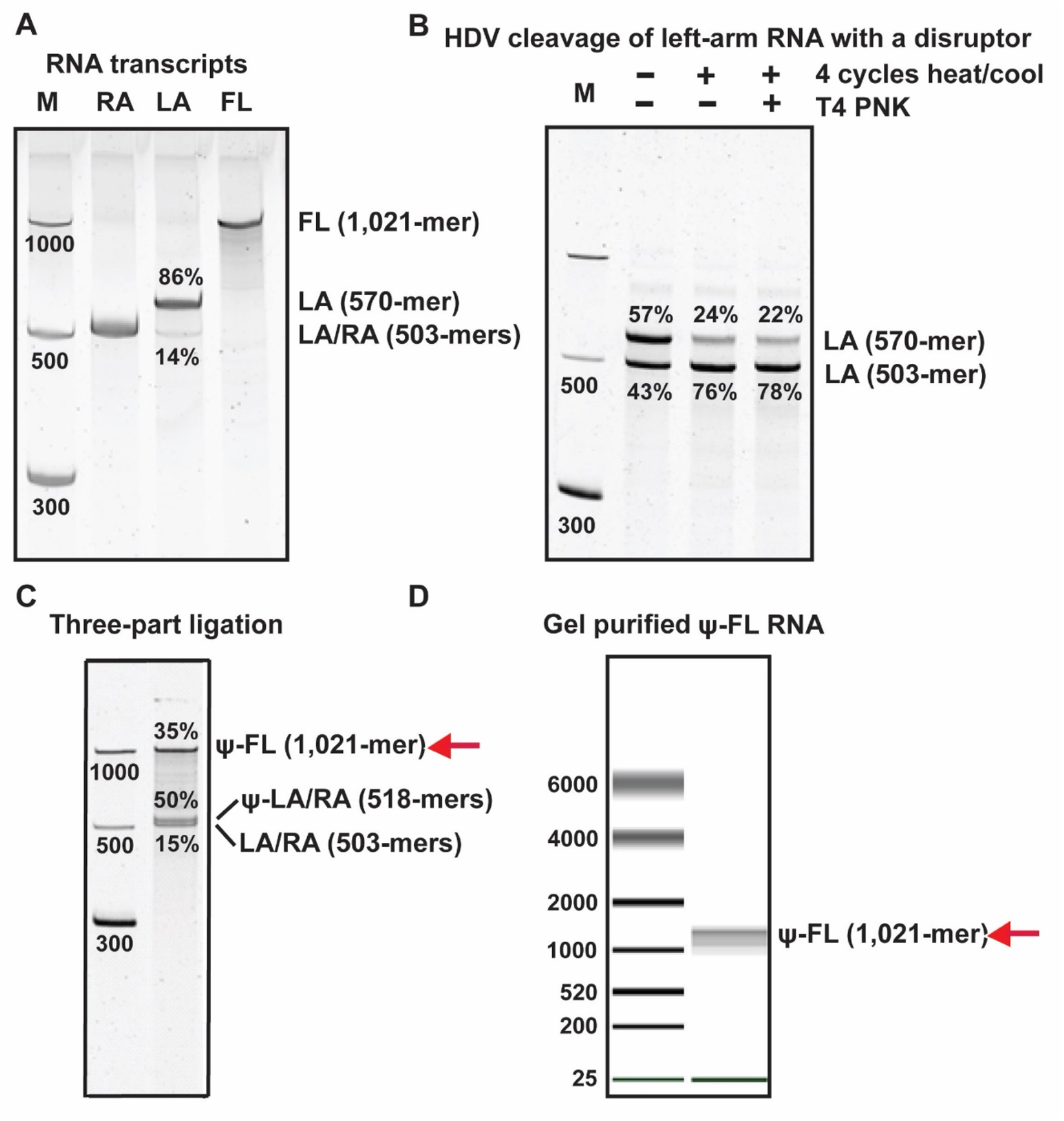
Step-by-step assembly of the 1 kb *PSMB2* RNA containing an internal ψ. (A) Denaturing PAGE (6%) analysis of the *in vitro* transcription products of the left-arm and right-arm RNA, showing that while the right-arm RNA migrated as a homogeneous 503-mer, the left-arm RNA migrated as a distribution of the 570-mer (86%) and 503-mer (14%). M: molecular weight markers; FL: full-length RNA (1 kb) made by *in vitro* transcription; LA: left-arm RNA; RA: right-arm RNA. (B) Denaturing PAGE (6%) analysis of HDV processing of the transcribed left-arm RNA over 4 heat-cool cycles, followed by T4 PNK hydrolysis. (C) Denaturing PAGE (6%) analysis of a 3-part splint ligation, yielding 35% as the ψ-containing full-length RNA (ψ-FL, 1 kb), 50% as the mixture of 2-part ligation products consisting of the left-arm and the right-arm each joined with the ψ-containing RNA [(LA-ψ) + (ψ-RA)] (518-mer each), and 15% as the mixture of un-ligated left-arm and right-arm RNA (503-mer each). Denaturing PAGE (6%) gels in panels A-C were each stained by SYBR-Gold. (D) Bioanalyzer (Agilent) capillary gel analysis of the 1 kb ψ-containing *PSMB2* RNA purified from a 1.2% agarose gel.

**Figure 5.**
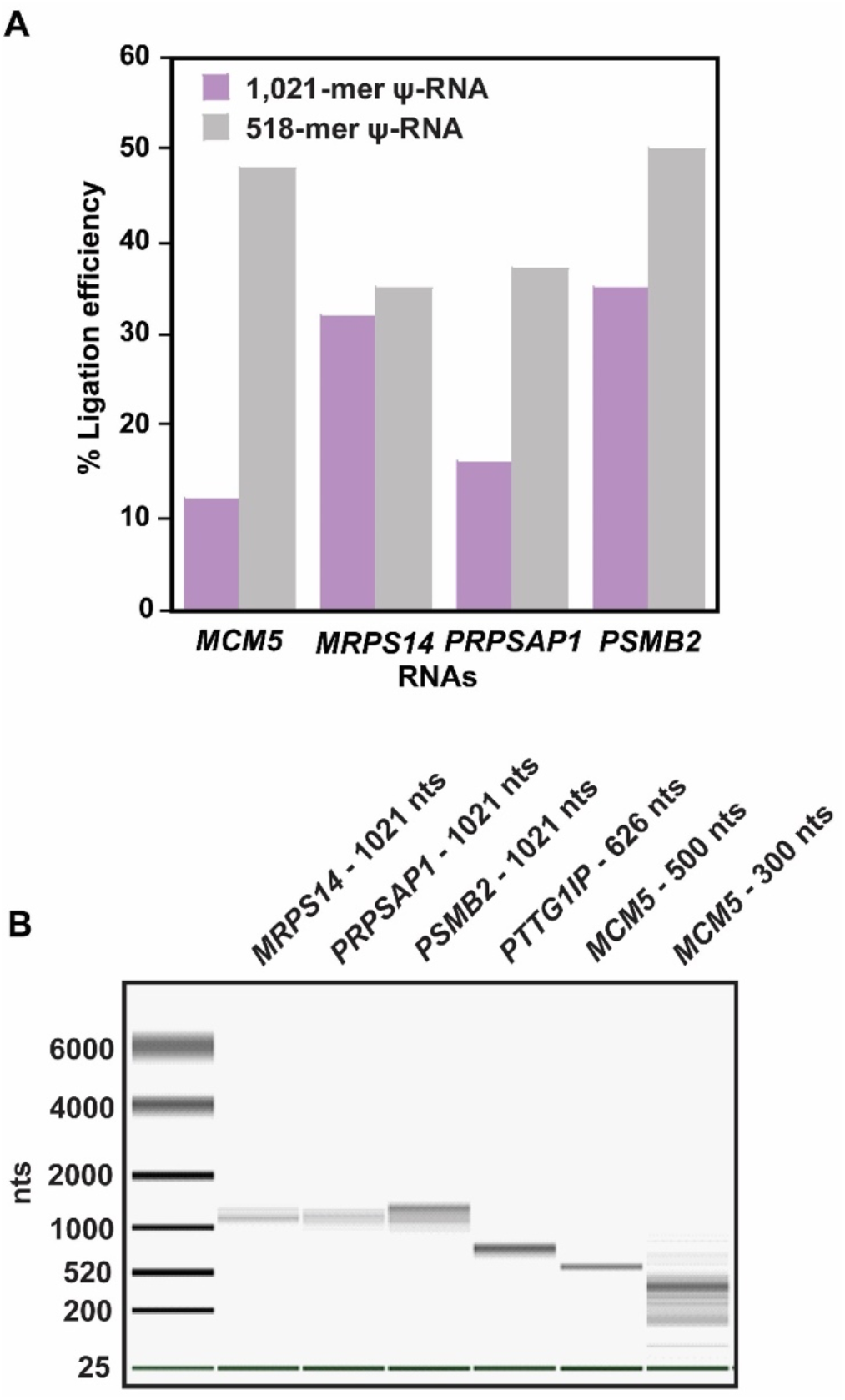
Efficiencies of 2-part and 3-part joining in a 3-part splint ligation reaction. (A) Efficiency of ligation by 2-part joining of the left-arm or right-arm RNA with a 15-mer to synthesize the 518-mer ψ-RNA is shown in grey, while that by 3-part joining of all three RNAs to synthesize the 1 kb ψ-RNA is shown in purple. (B) The quality of gel-purified ψ-containing long RNA by a capillary gel analysis. *MRPS14* RNA, *PRPSAP1* RNA, and the *PSMB2* RNA, all of 1,021 nts, were assembled from a left-arm (503-mer), a right-arm (503-mer), and a 15-mer ψ-RNA. *PTTG1IP* RNA of 626 nts was assembled from a left-arm (503-mer), a right-arm (108-mer), and a 15-mer ψ-RNA; *MCM5* RNA of 300 nts was assembled from a left-arm (141-mer), a right-arm (144-mer), and a 15-mer ψ-RNA; while *MCM5* RNA of 500 nts was assembled from a left-arm (242-mer), a right-arm (243-mer), and a 15-mer ψ-RNA.

### Optimization of splint ligation

We optimized two parameters of the splint ligation reaction. The first is the concentration of the DNA disruptor relative to its complementary RNA, which can strongly influence the efficiency of an inter-molecular ligation reaction. Using the *PSMB2* mRNA as an example, we designed a 2-part splint ligation reaction, in which the left- and right-arm RNA, each hybridized to a DNA disrupter, were aligned on a 24-mer DNA splint. We monitored the ligation efficiency as a function of the concentration of each disruptor relative to its complementary RNA (Supplementary Figure S1A). Specifically, the left- and right-arm disruptor were mixed in equal concentration, the left- and right-arm RNA were mixed in equal concentration, while the molar ratio of each disruptor to its RNA varied. Analysis of the molar ratio of the left-arm disruptor to the left-arm RNA as an indicator revealed no ligation in the absence of the disruptor, supporting the importance of the disruptor for ligation of long RNAs. In contrast, increasing concentration of the disruptor increased the ligation efficiency until the molar ratio reached ~18.0 at the start of a plateau. This molar ratio could vary with the length of the long RNA. In our example of joining a 500-mer left-arm RNA with a 500-mer right-arm RNA, we use 10 μM of each disruptor to 0.4 μM of the RNA in a molar ratio of 25, which is more than sufficient to reach the plateau of ligation efficiency.

The second parameter is the temperature and time of splint ligation. Given the conformational heterogeneity of long RNAs, the accessibility of each to hybridize to the disruptor may be discriminated. RNA secondary and tertiary structures have been proposed as a main contributor to ligation bias (57,58). Using the *PSMB2* mRNA in a 2-part splint ligation as above (Supplementary Figure S1B), we observed a progressive increase in the ligation efficiency with increasing temperature from 16 to 25 to 37 °C, indicating that higher temperatures help to unwind internal structures of long RNAs to facilitate ligation. At each temperature, we observed a plateau of the ligation efficiency starting at ~20 min, indicating that this is the time required to assemble the active pre-ligation complex. The consistency of the time across all three temperatures indicates that, once the temperature-dependent formation of the active pre-ligation complex is established, T4 RNL2 readily catalyzes ligation. Indeed, the intrinsic catalytic efficiency of T4 RNL2 is on the time scale of seconds (59). Thus, for ligation of long RNAs, temperature is the driving force to form the pre-ligation complex, which is a slow step and is followed by a fast step that catalyzes ligation. The identified time of 20 min is shorter than the commonly recommended time (> 1h) of splint ligation (60). The shorter time provides an option to reduce RNA degradation during a longer incubation time. Notably, we also observed a slow and gradual increase of ligation efficiency over a time scale of hours (not shown), indicating the possibility of rearrangement of the left-arm and right-arm RNA to make additional ends accessible for ligation.

### Assembly and purification of 1 kb RNA containing a site-specific internal ψ

We provide a step-by-step procedure to assemble a 1 kb-long RNA containing ψ at its natural sequence context. Using *PSMB2* as an example, we show that while the left-arm and right-arm RNA were both generated by *in vitro* transcription, and while the right-arm RNA migrated as a homogeneous 503-mer (the transcribed length), the left-arm RNA displayed a distribution between 86% as a 570-mer (the transcribed length) and 14% as a 503-mer (the HDV-cleaved length) (Figure 4A). Thus, a low level of the HDV cleavage reaction had occurred during *in vitro* transcription. This cleavage reaction was further activated upon repeated heat-cool cycles in the presence of the left-arm disruptor, generating 78% of the cleaved left-arm RNA whose 2’,3’-cyclic phosphate at the 3’-end was then removed by T4 PNK (Figure 4B). The 3’-end processed leftarm RNA was ligated with the right-arm RNA, together with the 15-mer ψ-containing synthetic RNA, in a 3-part splint ligation. Analysis of a 6% denaturing PAGE (Figure 4C) allowed quantification of each ligation product as the fraction of the input RNA. We found 35% as the 3-part ligation product of 1 kb ψ-containing RNA, 50% as the 2-part 518-mer ligation products, representing a mixture of the left-arm and right-arm RNA each ligated to the 15-mer, and 15% as a mixture of the un-ligated left-arm and right-arm 503-mer RNA. Notably, the yield of the 1 kb RNA (35%) is 3-5-fold higher than the reported yields (7-15%) of RNA of similar length generated by a 3-part splint ligation that included disruptors or a long splint but lacked ribozyme-processing of the left-arm RNA (30,37). Thus, improving the 3’-end homogeneity of the left-arm RNA is the major determinant of the higher yield.

The improved ligation yields afforded purification of the 1 kb ψ-containing RNA from other RNAs of the ligation reaction. While the 1 kb RNA migrated to a distinct position in a denaturing PAGE, we recovered little from the gel by extraction, consistent with the notion that RNA of > 600 nts is difficult to extract from denaturing PAGE (61). Instead, we found that the 1 kb RNA, which also migrated to a distinct position in an agarose gel, can be extracted with a yield of 40-50% and purified by a Zymo cartridge, leading to a product that showed a single band on an Agilent Bioanalyzer gel (Figure 4D). This extraction-purification method is more suitable for long RNA than multiple rounds of electroelution, which is prone to degradation of RNA (37).

### Ligation efficiency dependent on the length and sequence context

We quantified the ligation efficiency by denaturing PAGE. Among different sequences of synthetic RNAs, the efficiencies of 3-part ligation to generate the 1 kb RNA ranged in 10-35%, while that of 2-part ligation to generate a mixture of the 518-mer RNAs (e.g., Figure 4C) ranged in 35-53% (Figure 5A). Thus, the efficiency varies depending on the sequence context in both the 3-part and 2-part ligation reactions. For each RNA, however, the efficiency is consistently higher in 2-part ligation than in 3-part ligation, although the difference between the two also varies depending on the sequence context. These results emphasize the importance of the sequence context in ligation efficiency.

Several synthetic ψ-containing *MCM5* RNAs of different lengths were prepared by 3-part ligation. As the length decreased from 1,021- to 500- and to 300-mer, the yield of the fully ligated product increased from 10 to 14 to 18%. This trend is consistent with the notion that shorter length decreases conformational heterogeneity of the RNA to facilitate ligation. Analysis of the gel-purified RNAs on an Agilent Bioanalyzer (Figure 5B) showed homogeneity of the 1 kb and 500-mer RNAs as purified from an agarose gel. While this was not the case for the 300-mer (last lane), we found that switching to PAGE purification afforded purification to homogeneity for this shorter length (not shown).

### Sequencing analysis across ligation junctions

Errors in 3-part splint ligation cannot be excluded, given the heterogeneity issues of kb-long RNA. Although kb-long RNAs have been evaluated for fidelity as a template for cellular protein synthesis (37), this is an ensemble analysis that does not determine the fraction of the template that is assembled correctly or incorrectly. At the single-molecule level, while Illumina sequencing was used to determine the sequence accuracy of ligation in a 3-part splint reaction, the accuracy was only examined for the left-junction, but not for the right-junction (37). It was found that a small fraction of the ligated sequence at the left junction had incorrect nucleotides (37). These issues could be due to the production of short reads by Illumina sequencing. We therefore used nanopore sequencing to report long reads across both ligation junctions in each kb-long RNA generated by our 3-part splint ligation.

The results showed that, across all 5 ψ-mRNAs, the sequence at each ligation junction located 7 nts from either site of the modification, is homogeneous and accurate (Figure 6). The lack of even a trace of mismatches across ligation of distinct sequence contexts among the 5 ψ-mRNAs is remarkable. Thus, the splint ligation in our optimized condition ensured ligation in the correct order and sequence at both the 5’- and 3’-end. Additionally, we observed the U-to-C mismatch corresponding to the ψ in each mRNA, supporting the notion that nanopore reads the modification as a basecalling error. The varying frequencies of the U-to-C mismatch among the 5 ψ-mRNAs indicate differences in the nanopore detection of each modification in different sequence contexts. Notably, we also detected other mismatches adjacent to the ψ in some of these mRNAs, which were not present in the respective IVT control, indicating that they are errors specific to nanopore sequencing. Most likely, they are errors induced by the presence of ψ. Overall, this work demonstrates that our 3-part splint ligation produces kb-long RNA with precise sequence at each ligation junction, providing confidence for subsequent investigation of the ligated RNA.

**Figure 6.**
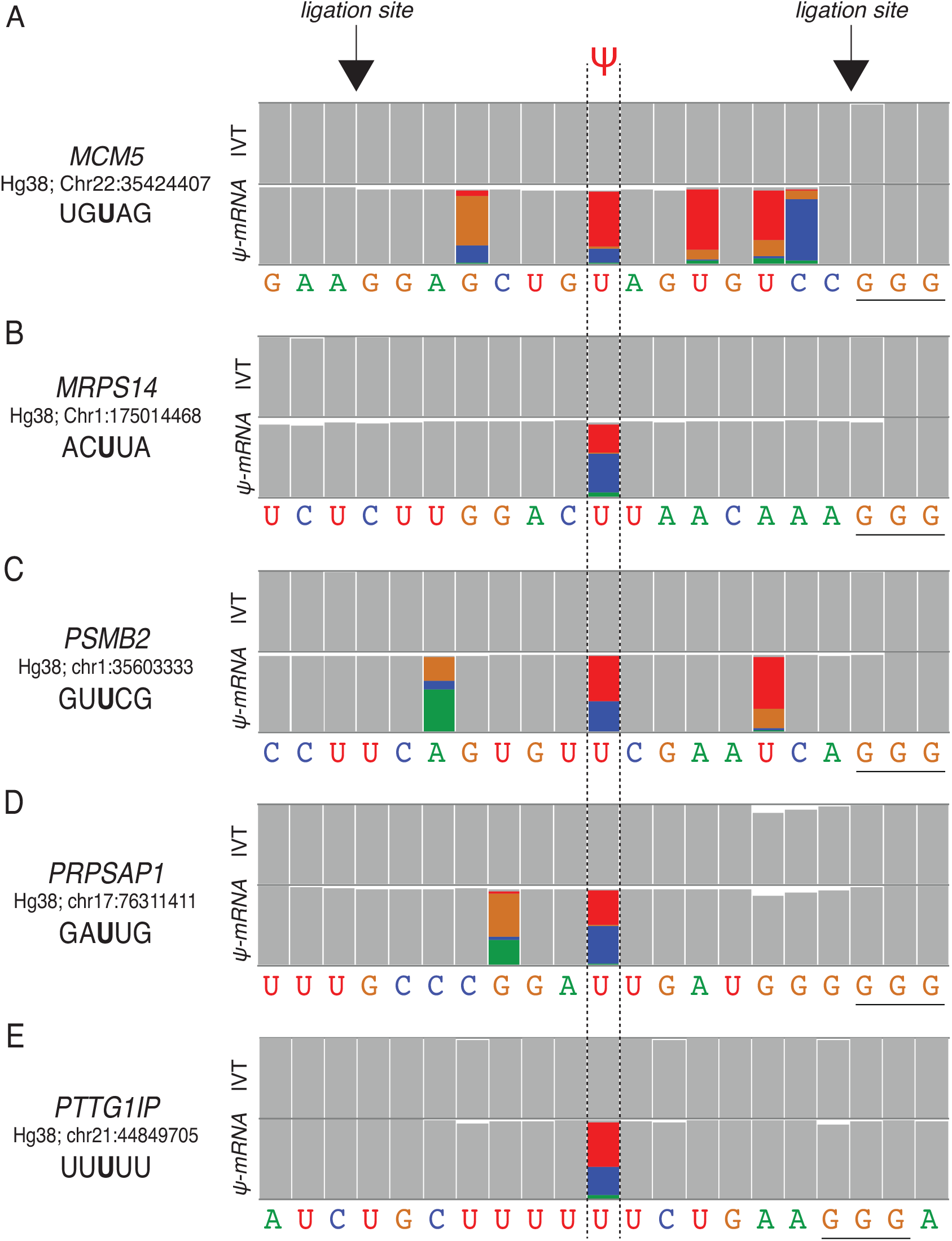
Nanopore sequencing across ligation junctions of ψ-mRNAs generated by 3-part splint ligation. Sequencing reads of ψ-mRNA of (A) *MCM5*, (B) *MRPS14*, (C) *PSMB2*, (D) *PRPSAP1*, and (E) *PTTG1IP* are shown by a representative snapshot from the integrated genome viewer (IGV) of aligned nanopore reads to the hg38 genome (GRCh38 p10) at previously annotated ψ sites. The sequence for each mRNA is shown below, where nucleotides A is shown in green, C in blue, G in brown, and U in red. Highlighted are miscalled bases of each mRNA, while grey indicates corrected called bases. The ligation junctions are marked by arrows, showing homogeneous and accurate sequences for each mRNA. The GGG immediately following the ligation site of the right-arm RNA is underlined, representing the initiation site of T7 transcription of the right-arm RNA. Except for the ψ-mRNA for *PTTG1Ip*, which was generated as a 600-mer, all others were generated as a 1 kb-mer.

## DISCUSSION

Synthesis of kb-long RNA by 3-part splint ligation has historically produced low yields (< 2%). Inclusion of DNA disruptors proximal to the ligation sites, or using a long splint DNA, has increased the yield (to 5-15%) (30,37). We report here a further improvement of the yield (to 10-35%) by using a *cis*-acting HDV ribozyme to generate the left-arm RNA with a homogeneous 3’-end in the presence of DNA disruptors. We find that the yield varies with the RNA length and sequence context, suggesting that optimization of each ligation is necessary. In some cases, we find that increasing the splint from a 39-mer to a 50-mer improves the yield. While increasing the molar ratio of the disruptors was previously reported to improve the yield (30), we find no improvement in our examples. Nor do we find improvement in our examples by varying the heat-cool temperatures for ribozyme cleavage prior to ligation, although we do find improvement by elevating the temperature of the ligation reaction itself. Thus, each sequence and length for ligation is distinct, and that several factors should be considered to achieve the maximum yield of ligation. These factors include the length of the DNA splint, molar ratio of disruptors to the left- and right-arm RNAs, and temperatures for the ribozyme cleavage and for the RNL2-catalyzed ligation.

A major step forward of this work is the ability to purify kb-long RNA with reasonable yields for nanopore sequencing analysis of the full-length RNA. Notably, RNA longer than hundreds of nucleotides is extremely difficult to isolate and purify, usually appearing as a smear in denaturing PAGE. This is due to the propensity of long RNA to degradation and to the inherent structural heterogeneity that leads to a population of transient isoforms that change continuously and dynamically in gel analysis. Importantly, we find that gel extraction, followed by purification through a Zymo cartridge, is the best method for isolation of kb-long RNA than alternative methods using an affinity tag (see below). Specifically, we find that the full-length RNA is most productively extracted from an agarose gel (1.2%), rather than a denaturing PAGE (6%). Because the yield of gel extraction is typically 50% and the yield through a Zymo cartridge is nearly stoichiometric, we use this estimation as a guide to design the amounts of input RNAs in each 3-part ligation reaction.

We have attempted different approaches to isolate kb-long RNA using an affinity tag but found that none produced the yield as high as extraction from an agarose gel. We describe our approaches for consideration by others interested in working with long RNA. (i) We have prepared the DNA splint with a biotin tag and used it to capture the ligated RNAs by a streptavidin resin. The bound RNAs were released from the resin by heat and analyzed on a denaturing PAGE. While the purity of the full-length RNA increased by 2-fold, only 10% of it was recovered. (ii) We have tested a two-step affinity-hybridization protocol, where we used one biotinylated probe for the left-arm to purify left arm-containing RNAs (products of both 2-part and 3-part ligation), followed by using a second biotinylated probe for the right-arm to purify right arm-containing fulllength RNA. We recovered only 1-2% of the full-length RNA. (iii) We have considered adding a poly(rA) tail to the right-arm RNA, which after ligation could be purified by a biotinylated oligo(dT) probe. However, this method would also pull down un-ligated and 2-part ligated right-arm RNA. Although none of the three affinity-based purification methods was satisfactory, we provide details in the Materials and Methods section as a basis for future improvement.

Our improved yields of 3-part splint ligation, combined with isolation of kb-long RNA, have enabled us to verify sequence accuracy across ligation junctions by nanopore sequencing. This demonstrates the ability of our method to generate long RNAs not only as synthetic standards for nanopore sequencing, but also as reagents for broader applications in RNA research and RNA-based therapies. Notably, short RNAs with an internal modification have proven valuable for RNA research to investigate RNA conformational dynamics and protein interacting partners (62–66). Long RNAs with a site-specific modification can now be used in cellular protein synthesis to determine the effect of the modification. Indeed, RNA biology frequently involves long RNA, such as excision of an intron (average of 5 kb), folding of rRNA (4-5 kb of the 28S and 1.9 kb of the human 18S), and regulation of gene expression by long non-coding RNAs (1-10 kb). In each case, a long RNA can be synthesized as a probe containing an internal fluorophore, or a pair of fluorophores, that respond to environmental changes and undergo fluorescence resonance energy transfer. We envision that our method will pave the way for a better understanding of each of these processes.

## DATA AVAILABILITY

Supplementary data are available at NAR Online.

## ACKNOWLEDGEMENTS

We thank all members of the Hou lab for discussion.

## REFERENCES

1. Wiener, D. and Schwartz, S. (2021) The epitranscriptome beyond m(6)A. Nat Rev Genet, 22, 119–131.

2. Dixit, S. and Jaffrey, S.R. (2022) Expanding the epitranscriptome: Dihydrouridine in mRNA. PLoS Biol, 20, e3001720.

3. Roundtree, I.A., Evans, M.E., Pan, T. and He, C. (2017) Dynamic RNA Modifications in Gene Expression Regulation. Cell, 169, 1187–1200.

4. Delaunay, S. and Frye, M. (2019) RNA modifications regulating cell fate in cancer. Nat Cell Biol, 21, 552–559.

5. Jonkhout, N., Tran, J., Smith, M.A., Schonrock, N., Mattick, J.S. and Novoa, E.M. (2017) The RNA modification landscape in human disease. RNA, 23, 1754–1769.

6. Li, X., Zhu, P., Ma, S., Song, J., Bai, J., Sun, F. and Yi, C. (2015) Chemical pulldown reveals dynamic pseudouridylation of the mammalian transcriptome. Nat Chem Biol, 11, 592–597.

7. Garalde, D.R., Snell, E.A., Jachimowicz, D., Sipos, B., Lloyd, J.H., Bruce, M., Pantic, N., Admassu, T., James, P., Warland, A. et al. (2018) Highly parallel direct RNA sequencing on an array of nanopores. Nat Methods, 15, 201–206.

8. Workman, R.E., Tang, A.D., Tang, P.S., Jain, M., Tyson, J.R., Razaghi, R., Zuzarte, P.C., Gilpatrick, T., Payne, A., Quick, J. et al. (2019) Nanopore native RNA sequencing of a human poly(A) transcriptome. Nat Methods, 16, 1297–1305.

9. Liu, H., Begik, O., Lucas, M.C., Ramirez, J.M., Mason, C.E., Wiener, D., Schwartz, S., Mattick, J.S., Smith, M.A. and Novoa, E.M. (2019) Accurate detection of m(6)A RNA modifications in native RNA sequences. Nat Commun, 10, 4079.

10. Begik, O., Lucas, M.C., Pryszcz, L.P., Ramirez, J.M., Medina, R., Milenkovic, I., Cruciani, S., Liu, H., Vieira, H.G.S., Sas-Chen, A. et al. (2021) Quantitative profiling of pseudouridylation dynamics in native RNAs with nanopore sequencing. Nat Biotechnol, 39, 1278–1291.

11. Huang, S., Zhang, W., Katanski, C.D., Dersh, D., Dai, Q., Lolans, K., Yewdell, J., Eren, A.M. and Pan, T. (2021) Interferon inducible pseudouridine modification in human mRNA by quantitative nanopore profiling. Genome Biol, 22, 330.

12. Fleming, A.M., Mathewson, N.J., Howpay Manage, S.A. and Burrows, C.J. (2021) Nanopore Dwell Time Analysis Permits Sequencing and Conformational Assignment of Pseudouridine in SARS-CoV-2. ACS Cent Sci, 7, 1707–1717.

13. Parker, M.T., Knop, K., Sherwood, A.V., Schurch, N.J., Mackinnon, K., Gould, P.D., Hall, A.J., Barton, G.J. and Simpson, G.G. (2020) Nanopore direct RNA sequencing maps the complexity of Arabidopsis mRNA processing and m(6)A modification. Elife, 9.

14. Price, A.M., Hayer, K.E., McIntyre, A.B.R., Gokhale, N.S., Abebe, J.S., Della Fera, A.N., Mason, C.E., Horner, S.M., Wilson, A.C., Depledge, D.P. et al. (2020) Direct RNA sequencing reveals m(6)A modifications on adenovirus RNA are necessary for efficient splicing. Nat Commun, 11, 6016.

15. Lorenz, D.A., Sathe, S., Einstein, J.M. and Yeo, G.W. (2020) Direct RNA sequencing enables m(6)A detection in endogenous transcript isoforms at base-specific resolution. RNA, 26, 19–28.

16. Pyle, A.M. (2008) Translocation and unwinding mechanisms of RNA and DNA helicases. Annu Rev Biophys, 37, 317–336.

17. Barbier, A.J., Jiang, A.Y., Zhang, P., Wooster, R. and Anderson, D.G. (2022) The clinical progress of mRNA vaccines and immunotherapies. Nat Biotechnol, 40, 840–854.

18. Kariko, K., Whitehead, K. and van der Meel, R. (2021) What does the success of mRNA vaccines tell us about the future of biological therapeutics? Cell Syst, 12, 757–758.

19. Musier-Forsyth, K., Usman, N., Scaringe, S., Doudna, J., Green, R. and Schimmel, P. (1991) Specificity for aminoacylation of an RNA helix: an unpaired, exocyclic amino group in the minor groove. Science, 253, 784–786.

20. Beuning, P.J., Yang, F., Schimmel, P. and Musier-Forsyth, K. (1997) Specific atomic groups and RNA helix geometry in acceptor stem recognition by a tRNA synthetase. Proc Natl Acad Sci U S A, 94, 10150–10154.

21. Musier-Forsyth, K. and Schimmel, P. (1992) Functional contacts of a transfer RNA synthetase with 2’-hydroxyl groups in the RNA minor groove. Nature, 357, 513–515.

22. Musier-Forsyth, K. and Schimmel, P. (1994) Acceptor helix interactions in a class II tRNA synthetase: photoaffinity cross-linking of an RNA miniduplex substrate. Biochemistry, 33, 773–779.

23. Sakaguchi, R., Giessing, A., Dai, Q., Lahoud, G., Liutkeviciute, Z., Klimasauskas, S., Piccirilli, J., Kirpekar, F. and Hou, Y.M. (2012) Recognition of guanosine by dissimilar tRNA methyltransferases. RNA, 18, 1687–1701.

24. Silber, R., Malathi, V.G. and Hurwitz, J. (1972) Purification and properties of bacteriophage T4-induced RNA ligase. Proc Natl Acad Sci U S A, 69, 3009–3013.

25. Nandakumar, J., Shuman, S. and Lima, C.D. (2006) RNA ligase structures reveal the basis for RNA specificity and conformational changes that drive ligation forward. Cell, 127, 71–84.

26. Ho, C.K. and Shuman, S. (2002) Bacteriophage T4 RNA ligase 2 (gp24.1) exemplifies a family of RNA ligases found in all phylogenetic domains. Proc Natl Acad Sci U S A, 99, 12709–12714.

27. Stark, M.R., Pleiss, J.A., Deras, M., Scaringe, S.A. and Rader, S.D. (2006) An RNA ligase-mediated method for the efficient creation of large, synthetic RNAs. RNA, 12, 2014–2019.

28. Keyhani, S., Goldau, T., Blumler, A., Heckel, A. and Schwalbe, H. (2018) Chemo-Enzymatic Synthesis of Position-Specifically Modified RNA for Biophysical Studies including Light Control and NMR Spectroscopy. Angew Chem Int Ed Engl, 57, 12017–12021.

29. Hoffmann, P.U. and McLaughlin, L.W. (1987) Synthesis and reactivity of intermediates formed in the T4 RNA ligase reaction. Nucleic Acids Res, 15, 5289–5303.

30. Zhovmer, A. and Qu, X. (2016) Proximal disruptor aided ligation (ProDAL) of kilobase-long RNAs. RNA Biol, 13, 613–621.

31. Chu, W.C. and Horowitz, J. (1989) 19F NMR of 5-fluorouracil-substituted transfer RNA transcribed in vitro: resonance assignment of fluorouracil-guanine base pairs. Nucleic Acids Res, 17, 7241–7252.

32. Cazenave, C. and Uhlenbeck, O.C. (1994) RNA template-directed RNA synthesis by T7 RNA polymerase. Proc Natl Acad Sci U S A, 91, 6972–6976.

33. Yin, Y. and Carter, C.W., Jr. (1996) Incomplete factorial and response surface methods in experimental design: yield optimization of tRNA(Trp) from in vitro T7 RNA polymerase transcription. Nucleic Acids Res, 24, 1279–1286.

34. Kholod, N., Vassilenko, K., Shlyapnikov, M., Ksenzenko, V. and Kisselev, L. (1998) Preparation of active tRNA gene transcripts devoid of 3’-extended products and dimers. Nucleic Acids Res, 26, 2500–2501.

35. Schurer, H., Lang, K., Schuster, J. and Morl, M. (2002) A universal method to produce in vitro transcripts with homogeneous 3’ ends. Nucleic Acids Res, 30, e56.

36. Strobel, S.A. and Cech, T.R. (1993) Tertiary interactions with the internal guide sequence mediate docking of the P1 helix into the catalytic core of the Tetrahymena ribozyme. Biochemistry, 32, 13593–13604.

37. Hertler, J., Slama, K., Schober, B., Ozrendeci, Z., Marchand, V., Motorin, Y. and Helm, M. (2022) Synthesis of point-modified mRNA. Nucleic Acids Res.

38. Chowrira, B.M., Pavco, P.A. and McSwiggen, J.A. (1994) In vitro and in vivo comparison of hammerhead, hairpin, and hepatitis delta virus self-processing ribozyme cassettes. J Biol Chem, 269, 25856–25864.

39. Been, M.D., Perrotta, A.T. and Rosenstein, S.P. (1992) Secondary structure of the selfcleaving RNA of hepatitis delta virus: applications to catalytic RNA design. Biochemistry, 31, 11843–11852.

40. Golden, B.L. (2011) Two distinct catalytic strategies in the hepatitis delta virus ribozyme cleavage reaction. Biochemistry, 50, 9424–9433.

41. Masek, T., Vopalensky, V., Suchomelova, P. and Pospisek, M. (2005) Denaturing RNA electrophoresis in TAE agarose gels. Anal Biochem, 336, 46–50.

42. Tavakoli, S., Nabizadehmashhadtoroghi, M., Makhamreh, A., Gamper, H., Razapour, N.K., Hou, Y.M., Wanunu, M. and Rouhanifard, S.H. (2022) Semi-quantitative detection of pseudouridine modifications and type I/II hypermodifications in human mRNAs using direct and long-read sequencing. Nat Commun.

43. Martin, C.T. and Coleman, J.E. (1989) T7 RNA polymerase does not interact with the 5’-phosphate of the initiating nucleotide. Biochemistry, 28, 2760–2762.

44. Been, M.D. (1994) Cis-and trans-acting ribozymes from a human pathogen, hepatitis delta virus. Trends Biochem Sci, 19, 251–256.

45. Perrotta, A.T. and Been, M.D. (1992) Cleavage of oligoribonucleotides by a ribozyme derived from the hepatitis delta virus RNA sequence. Biochemistry, 31, 16–21.

46. Anderson, B.R., Muramatsu, H., Jha, B.K., Silverman, R.H., Weissman, D. and Kariko, K. (2011) Nucleoside modifications in RNA limit activation of 2’-5’-oligoadenylate synthetase and increase resistance to cleavage by RNase L. Nucleic Acids Res, 39, 9329–9338.

47. Kariko, K., Buckstein, M., Ni, H. and Weissman, D. (2005) Suppression of RNA recognition by Toll-like receptors: the impact of nucleoside modification and the evolutionary origin of RNA. Immunity, 23, 165–175.

48. Anderson, B.R., Muramatsu, H., Nallagatla, S.R., Bevilacqua, P.C., Sansing, L.H., Weissman, D. and Kariko, K. (2010) Incorporation of pseudouridine into mRNA enhances translation by diminishing PKR activation. Nucleic Acids Res, 38, 5884–5892.

49. Eyler, D.E., Franco, M.K., Batool, Z., Wu, M.Z., Dubuke, M.L., Dobosz-Bartoszek, M., Jones, J.D., Polikanov, Y.S., Roy, B. and Koutmou, K.S. (2019) Pseudouridinylation of mRNA coding sequences alters translation. Proc Natl Acad Sci U S A, 116, 23068–23074.

50. Carlile, T.M., Rojas-Duran, M.F., Zinshteyn, B., Shin, H., Bartoli, K.M. and Gilbert, W.V. (2014) Pseudouridine profiling reveals regulated mRNA pseudouridylation in yeast and human cells. Nature, 515, 143–146.

51. Khoddami, V., Yerra, A., Mosbruger, T.L., Fleming, A.M., Burrows, C.J. and Cairns, B.R. (2019) Transcriptome-wide profiling of multiple RNA modifications simultaneously at single-base resolution. Proc Natl Acad Sci U S A, 116, 6784–6789.

52. Safra, M., Nir, R., Farouq, D., Vainberg Slutskin, I. and Schwartz, S. (2017) TRUB1 is the predominant pseudouridine synthase acting on mammalian mRNA via a predictable and conserved code. Genome Res, 27, 393–406.

53. Smith, A.M., Jain, M., Mulroney, L., Garalde, D.R. and Akeson, M. (2019) Reading canonical and modified nucleobases in 16S ribosomal RNA using nanopore native RNA sequencing. PLoS One, 14, e0216709.

54. Schwartz, S., Bernstein, D.A., Mumbach, M.R., Jovanovic, M., Herbst, R.H., Leon-Ricardo, B.X., Engreitz, J.M., Guttman, M., Satija, R., Lander, E.S. et al. (2014) Transcriptome-wide mapping reveals widespread dynamic-regulated pseudouridylation of ncRNA and mRNA. Cell, 159, 148–162.

55. Makhamreh, A., Tavakoli, S., Gamper, H., Nabizadehmashhadtoroghi, N., Fallahi, A., Hou, Y.M., Rouhanifard, S.H. and Wanunu, M. (2022) Messenger-RNA modification standards and machine learning models facilitate absolute site-specific pseudouridine quantification. bioRxiv.

56. Ferre-D’Amare, A.R., Zhou, K. and Doudna, J.A. (1998) Crystal structure of a hepatitis delta virus ribozyme. Nature, 395, 567–574.

57. Hafner, M., Renwick, N., Brown, M., Mihailovic, A., Holoch, D., Lin, C., Pena, J.T., Nusbaum, J.D., Morozov, P., Ludwig, J. et al. (2011) RNA-ligase-dependent biases in miRNA representation in deep-sequenced small RNA cDNA libraries. RNA, 17, 1697–1712.

58. Zhuang, F., Fuchs, R.T., Sun, Z., Zheng, Y. and Robb, G.B. (2012) Structural bias in T4 RNA ligase-mediated 3’-adapter ligation. Nucleic Acids Res, 40, e54.

59. Chauleau, M. and Shuman, S. (2013) Kinetic mechanism of nick sealing by T4 RNA ligase 2 and effects of 3’-OH base mispairs and damaged base lesions. RNA, 19, 1840–1847.

60. Kershaw, C.J. and O’Keefe, R.T. (2012) Splint ligation of RNA with T4 DNA ligase. Methods Mol Biol, 941, 257–269.

61. Nilsen, T.W. (2013) Gel purification of RNA. Cold Spring Harb Protoc, 2013, 180–183.

62. Doudna, J.A., Szostak, J.W., Rich, A. and Usman, N. (1990) Chemical synthesis of oligoribonucleotides containing 2-aminopurine: substrates for the investigation of ribozyme function. J Org Chem, 55, 5547–5549.

63. Chow, C.S., Mahto, S.K. and Lamichhane, T.N. (2008) Combined Approaches to Site-Specific Modification of RNA. ACS Chem Biol, 3, 30–37.

64. Hobartner, C. and Micura, R. (2004) Chemical synthesis of selenium-modified oligoribonucleotides and their enzymatic ligation leading to an U6 SnRNA stem-loop segment. J Am Chem Soc, 126, 1141–1149.

65. Liu, W., Shin, D., Tor, Y. and Cooperman, B.S. (2013) Monitoring translation with modified mRNAs strategically labeled with isomorphic fluorescent guanosine mimetics. ACS Chem Biol, 8, 2017–2023.

66. Stone, M.D., Mihalusova, M., O’Connor C, M., Prathapam, R., Collins, K. and Zhuang, X. (2007) Stepwise protein-mediated RNA folding directs assembly of telomerase ribonucleoprotein. Nature, 446, 458–461.

