## Supplementary information for "Synthesis of Long RNA with a Site-Specific Modification by Enzymatic Splint Ligation"

**[1]Department of Biochemistry and Molecular Biology, Thomas Jefferson University, Philadelphia, PA, USA**

**[2]Department of Bioengineering, Northeastern University, Boston, MA, USA**

**[3]Department of Physics, Northeastern University, Boston, MA, USA**

**[*]Corresponding author:**

** (T) 215-503-4480; (F) 215-503-4954**

**ORCID: 0000-0001-6546-2597**

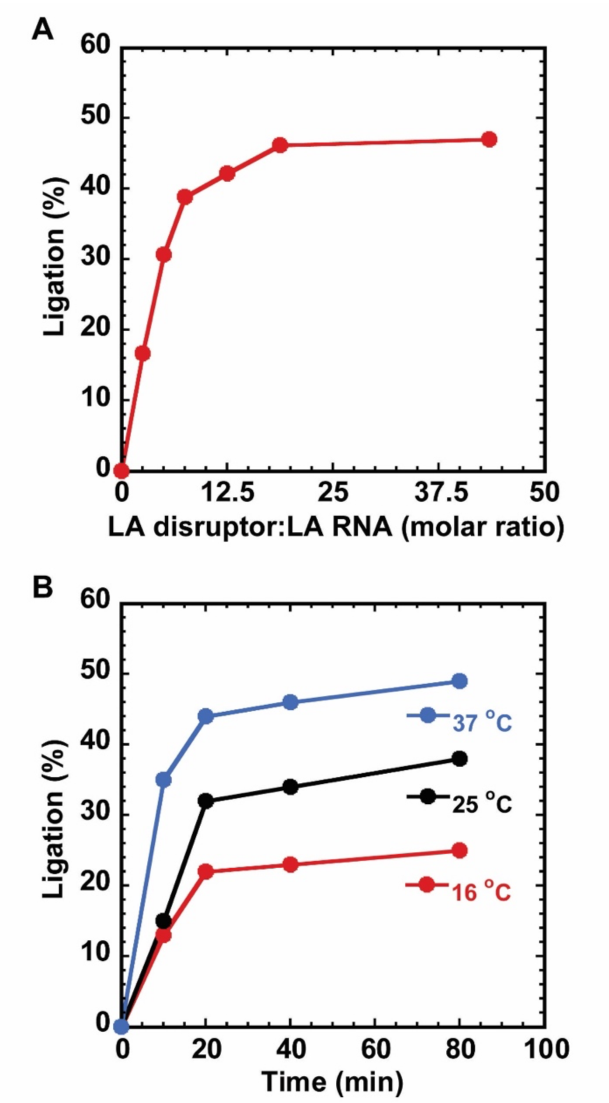


**Supplementary Figure S1**. Optimization of splint ligation. Assembly of a kb-long *PSMB2* mRNA by T4 RNL2-catalyzed ligation of a 500-mer left-arm RNA with a 500-mer right-arm RNA on a 12-mer DNA splint. (A) Efficiency of ligation (%) as a function of the molar ratio of the left-arm DNA disruptor relative to the left-arm RNA. (B) Efficiency of ligation (%) as a function of time achieved by T4 RNL2-catalyzed reaction at 16, 25, and 37 °C. The condition of ligation was as described in the standard 3-part ligation reaction.
